# Chronic subthreshold intermittent theta burst stimulation promotes structural axon initial segment plasticity in cortical neurons

**DOI:** 10.64898/2026.04.24.720748

**Authors:** Emily S King, Liz Jaeschke-Angi, Hakuei Fujiyama, Wicliffe Abraham, Jennifer Rodger, John NJ Reynolds, Darren Clarke, Jamie L Beros, Alexander D Tang

## Abstract

Repetitive transcranial magnetic stimulation (rTMS) is used widely in neuroscience to study and alter neural plasticity. The cellular mechanisms underlying the effect of rTMS on the brain remain unclear but is primarily thought to act via activity-dependent synaptic plasticity mechanisms. Here we investigated whether chronic repetitive magnetic stimulation *in vitro* and *in vivo* can induce another form of activity-dependent neural plasticity, axon initial segment (AIS) plasticity. Cortical neurons isolated from postnatal wild-type mice were stimulated with 6 hours of sham, repetitive magnetic stimulation in the form of intermittent theta-burst stimulation (iTBS), or 15 mM potassium chloride, with changes to AIS location and length measured +0 hours and +24 hours post-stimulation. In addition, adult transgenic mice expressing green fluorescent protein at the AIS received daily sham or iTBS over the primary motor cortices for 7 consecutive days and processed for microscopy 3 hours after the last stimulation. Analysis of neurons stimulated *in vitro* showed that chronic iTBS caused bidirectional and time-dependent shifts to the AIS position relative to the soma and a delayed shortening of the AIS length at +24 hours. In the adult mice, 7 consecutive days of daily iTBS decreased AIS lengths in layers 2/3 and 5 pyramidal neurons. Our findings provide *in vitro* and *in vivo* evidence that rTMS induces neuronal plasticity outside of the synapse, which may contribute to the long-lasting effect of rTMS on the brain with repeated stimulation protocols.

## Introduction

Neural plasticity is an essential process in the nervous system, underlying key neurological functions such as learning and memory, and is a common therapeutic target for many neurological conditions. As a result, there has been great interest in developing non-invasive methods that promote neural plasticity in the brain, such as repetitive transcranial magnetic stimulation (rTMS). While rTMS is most commonly used to alter neural activity to promote synaptic plasticity [1–6], experiments using rodent tissue have shown rTMS-induced changes to other activity-dependent plasticity mechanisms in neurons [7,8,6,9] and glia [5,10–12,6,9]. For example, stimulation with the intermittent theta burst stimulation (iTBS) protocol [13] has been shown to increase neuronal excitability through changes to intrinsic membrane properties [7–9], suggesting rTMS can alter neuronal compartments outside the synapse [14].

One neuronal compartment known to undergo activity dependent plasticity is the axon initial segment (AIS) [15–17], the site of action potential generation in mammalian neurons [18–22]. AIS plasticity is known to occur in the adult brain [17] in response to significant changes in neural activity, causing structural reorganisation of several scaffolding proteins at the AIS, including changes to AnkyrinG (AnkG) [17,23]. These structural changes manifest as a change to AIS position on the axon relative to the soma [15] or a change to AIS length [16,24,17,25,23,26]. The impact of shifting AIS position on action potential generation is complex, and depending on several factors such as soma size could either increase or decrease neuronal excitability [24]. However, changes to AIS length are more intuitive as AIS length is proportional to the number of ion channels involved in action potential generation and propagation, with decreased AIS lengths decreasing neural excitability and vice versa [15–17,27].

While AIS plasticity is an endogenous process [17], experimental studies using rodent tissue have also shown that it can be induced externally with brain stimulation. For example, increasing neural activity with bursts of optogenetic stimulation at “physiologically relevant” frequencies causes a distal shift of AIS position *in vitro* [15], and high frequency electrical stimulation of the perforant path *in vivo* induces long-term potentiation and a shortening of the AIS in dentate gyrus granule cells [25]. We recently extended this to non-invasive brain stimulation using static magnetic stimulation for 6 or 48 hours to promote structural AIS plasticity in cortical neurons *in vitro* [28]. However, changes to AIS plasticity with more widely used non-invasive techniques such as rTMS which are commonly delivered for days to weeks at a time in humans have not been investigated. Here we characterised the effect of repetitive magnetic stimulation in the form of chronic iTBS on AIS structural plasticity in cortical neurons. Given that iTBS is delivered to increase neural excitability and activity [13,29] and AIS plasticity serves as a homeostatic mechanism to stabilise neural activity [17], we hypothesised that chronic iTBS wiould induce structural AIS plasticity to reduce neural excitability (e.g., reduced AIS length). In primary cortical neurons isolated from postnatal mice, we provide proof of principle that 6 hours of iTBS induces acute and long-lasting AIS structural plasticity that is not due to neuronal injury. *In vivo,* stimulating the primary motor cortices of awake and freely moving adult mice with daily iTBS for 7 consecutive days caused a shortening of AIS lengths in layers 2/3 and 5 pyramidal neurons. Taken together, we provide *in vitro* and *in vivo* evidence that iTBS induces neuronal plasticity outside of the synapse, uncovering AIS plasticity as a novel neural plasticity mechanism recruited by repeated rTMS.

## Materials and methods

### Animals

All animal experimentation was approved by the University of Western Australia Animal Ethics Committee (AEC 03/100/1677 and 2022/ET000022) in accordance with the Australian code of practice for the care and use of animals for scientific purposes.

For *in vitro experiments*, wild-type postnatal day 1 C57Bl/6J mice of both sexes were purchased (Ozgene, Perth, Australia) and used on the day of delivery. For the *in vivo* stimulation experiments, we used a Cre-dependent transgenic mouse line that expresses green fluorescent protein (GFP) at the AIS (AnkG-GFP mice [23]). This transgenic line uses a FLeX cassette system to flip out the last exon of *Ank3* and replace it with the last exon of *Ank3* fused with the coding sequence for GFP, leading to GFP expression at the AIS after Cre-recombination. Mice of both sexes were used and group housed on a 12-hour light/dark cycle and given food and water *ad libitum*.

### Neuronal cell culture

Primary neuronal cell cultures were prepared using identical methods to [28]. Postnatal day 1 mice were euthanised with an overdose of sodium pentobarbitone (Lethabarb, Virbac) administered by an intraperitoneal injection. After confirmation of death, both cerebral cortices were dissected into serum free medium consisting of 2% B27 plus supplement (Gibco; Catalogue number A3582801), 0.25% GlutaMAX supplement (Gibco; Catalogue number 35050061) in Hibernate A medium (Gibco; Catalogue number A1247501). Following the removal of the meninges, dissected cortices were cut into smaller pieces (approx. 0.5 mm^2^) using a scalpel blade. Cortical tissue pieces were equilibrated at 30 °C for 10 min prior to incubation in filtered pre-warmed dissociation medium consisting of 2 mg/ml papain (Sigma-Aldrich; Catalogue number P4762) and 0.25% Glutamax (Gibco; Catalogue number 35050061) in Hibernate A medium without calcium and magnesium (BrainBits; Catalogue number Ha-Ca). This mixture was spun at 350 rpm at 30 °C for 30 min before the cortices were transferred into 2 ml of pre-warmed dissection medium. Cortices were triturated using a sterile glass Pasteur pipette (Corning) and the supernatant transferred into a centrifuge tube. After repeating the trituration process and supernatant removal for an additional two times, the collected supernatant was centrifuged at 12,000 rpm for 5 min. The supernatant was aspirated and the pellet was resuspended in culture medium consisting of 2% B27 plus supplement (Gibco), 0.25% Glutamax (Gibco) and 0.1% penicillin/streptomycin (Gibco; Catalogue number 15140163) in Neurobasal Plus medium (Gibco; Catalogue number A3582901). Cells were seeded at a density of 150,000 cells/coverslip onto 13 mm diameter coverslips in 24 well plates (Trajan) pre-coated with 50 µg/ml poly-d-lysine (Gibco; Catalogue number A3890401) and 40 µg/ml mouse type 1 laminin (Gibco; Catalogue number 23017015). Cell culture plates were incubated at 37 °C with 5% CO_2_ with half of the media replaced with fresh culture media at 3 and 7 days in vitro (DIV).

### *In vitro* stimulation

At DIV7, cultures were stimulated with sham, iTBS, or 15 mM KCl for 6 hours. KCl was chosen as a positive control as it has previously been shown to induce structural AIS plasticity *in vitro* [15,28]. Sham stimulated coverslips underwent a 50% media replacement before being placed in the incubator for 6 hours. For the KCl stimulated groups, KCl (Sigma Aldrich; Catalogue number P3911) was dissolved into the 50% media replacement and coverslips were placed into the incubator for 6 hours, a paradigm previously used by us and others to induce structural AIS plasticity *in vitro* [28,15]. For iTBS, a custom iron core circular coil (8 mm in height x 8 mm outer diameter) originally designed for rodent specific stimulation [30] was used to deliver the repetitive magnetic stimulation (rMS) inside the incubator. The coil sat 2 mm directly underneath the culture plates with the coverslips lying perpendicular to the coil windings. The iTBS protocol consisted of monophasic pulses of 10 trains of 3 pulses of 50 Hz stimulation repeated at 5Hz with an inter-train interval of 10s for 6 hours (total of 68,210 pulses) with a peak magnetic field intensity of 65 mT at the coil surface. This was the maximum magnetic field output that could be achieved for 6 hours in the miniaturised coils without altering coil temperature, avoiding heating of the cells and culture media in the wells. The stimulation was controlled by an arbitrary waveform generator (Agilent Technologies 335141B) connected to a bipolar voltage programmable power supply (KEPCO BOP 100-4M). To estimate the electric field induced in the neuronal cell layer, a simple 3D numerical approximation of Faraday’s law was modelled (MATLAB 2024B), accounting for the coil distance and the geometry of the coverslip and attenuation by the surrounding media. The neuronal layer was represented as a thin sheet laying on top of the glass coverslip, and the electric field distribution was visualised with smooth 3D surfaces and vector plots. The model computed an estimated peak induced electric field of 1.96V/m within the neuronal cell layer (Fig S.1).

At the end of stimulation, half of the coverslips were removed from the incubator and washed with 1X phosphate buffered saline (PBS pH 7.2 – 7.4) and fixed with 4% paraformaldehyde for 10 mins at room temperature to assess the immediate effects of stimulation (+0hrs group). The remaining coverslips received a 90% media replacement and were returned to the incubator until +24 hours post-stimulation, followed by a wash and fixation as described above (+24hrs group). For both groups, coverslips were stored in PBS with 0.1% sodium azide until further use.

### Pharmacological manipulation

A subset of coverslips were treated with tetrodotoxin (TTX – Abcam; Catalogue number ab120055) or mibefradil (Abcam: Catalogue number ab120343) to block voltage-gated sodium and L and T-type calcium channels respectively. Both blocking agents were made up to the manufacturer’s specifications and 1 hour prior to stimulation, blockers were added to the 50% media replacement to achieve a final concentration of 1 μM TTX and 3 μM of mibefradil. Coverslips then received control, KCl or iTBS as described above.

### *In vitro* immunofluorescence and microscopy

Coverslips were washed with three 10 min PBS washes followed by permeabilisation for 10 min with PBS containing 0.2% Triton-X. Coverslips were blocked for 1 hour in 10% normal goat serum and 0.2% Triton-X. Coverslips were incubated overnight at 4°C in PBS containing 0.2% Triton-X, 2% normal goat serum and primary antibodies for mouse-anti-Ankyrin G (1:600, Thermo Fisher Scientific; Catalogue number p33-8800 ab_2533145), chicken-anti-MAP2 (1:5000, Life Technologies; Catalogue number ab5392), and rabbit-anti-Glutamate decarboxylase 65-67 (GAD65-67) (1:1000, Life Technologies; Catalogue number ab36080). Following incubation, coverslips were washed with two 10 min PBS washes and incubated for 2 hours at room temperature in PBS containing 0.2% Triton-X, 2% normal goat serum with secondary antibodies goat-anti-mouse Alexa Fluor488 (1:600, Invitrogen), goat-anti-rabbit Alexa Fluor594 (1:600, Invitrogen) and goat-anti-chicken Alexa Fluor647 (1:600, Invitrogen). Coverslips were washed with two 10 min PBS washes and incubated in Hoechst dye for 10 min. For cell purity analysis, glial fibrillary acidic protein (GFAP) (anti-mouse 1:1000, Thermofisher; Catalogue number 14-9892-82) was labelled with MAP2 and Hoechst. A final PBS wash was applied before mounting the coverslips onto glass cover slides with Prolong Diamond Antifade Mountant (Life Technologies; catalogue number P36970). Coverslips were stored at 4 °C for a minimum of 48 hours prior to imaging.

A Nikon C2 confocal microscope (NIS Elements, 60x objective, oil immersion, NA=1.40, Nikon Plan Apo, Z step spacing of 0.125μm) using excitation and emission filters for DAPI (Hoescht), FITC (Alexa Fluro488), TRITC (Alexa Fluor594) and CY5 (Alexa Fluor647) was used to image the immunofluorescence. 15 fields of view (512 x 512 pixels) were captured evenly across each coverslip. For each image, 2X signal averaging was used to visualise Ankyrin G/AlexaFluor-488 channel. Maximum intensity Z-projections were generated in FIJI [31] and converted to RGB Tiff files before renaming to allow for blinded analysis. AIS lengths were measured using a custom MATLAB developed by Prof Matthew Grubb (Matthew Grubb (2020); freely available at https://www.mathworks.com/matlabcentral/fileexchange/28181-ais-quantification). In brief, the custom script quantifies Ankyrin G fluorescence along manually traced axons, defining the AIS start and end positions where the Ankyrin G signal crosses 33% of the maximum fluorescence [32,15,33]. Measurements of AIS distance from soma were done in FIJI by manually drawing a segmented line from the soma (identified by MAP2 staining) to the start position of the Ankyrin G labelling at the AIS. Neurons were excluded from the analysis if there was more than one AIS, a bifurcated AIS, or an AIS originating from a dendrite. Neurons were analysed separately depending on whether they were excitatory (GAD65-67 negative) or inhibitory (GAD65-67 positive). While quantitative analysis showed that our cell cultures contained both excitatory and inhibitory neurons, the majority were inhibitory neurons (Table S.1). For this reason, we restricted our analysis to primary cortical inhibitory neurons where meaningful statistical analysis could be made.

### Cell viability staining

The Image-IT^TM^ DEAD Green^TM^ Viability Stain (Invitrogen; Catalogue number 10291) assay was used to assess cell viability in a subset of coverslips. One microlitre of the stain was added per well at the end of the experiment and incubated for a further 30 mins. Coverslips were then washed with PBS, fixed with 4% PFA for 10 min at room temperature and stored in 1x PBS containing 0.1% sodium azide. Coverslips were processed for immunofluorescence to label MAP2 as described above and imaged on a Nikon C2 confocal microscope (20x objective, NA=0.75, Nikon Plan Apo, Z step spacing of 1 μm) using excitation and emission filters for DAPI (Hoescht), FITC (Alexa Fluro488), and CY5 (Alexa Fluor647). All images were renamed to allow for blinded analysis. Neurons were identified as dead or dying if they were MAP2^+ve^-Image-IT^+ve^-Hoechst^+ve^ whereas cells that were MAP2^-ve^-Image-IT^+ve^-Hoechst^+ve^ were identified as dead or dying non-neuronal cells.

### Stereotaxic surgery and adeno-associated virus (AAV) injections

11-week-old AnkG-GFP mice underwent stereotaxic surgery to deliver bilateral injections of AAV1-hSyn-Cre-P2A-TdTomato (Addgene; #107738), enabling the Cre-Lox recombination for GFP expression at the AIS. Mice received pre-operative analgesia (buprenorphine, 0.05 mg/kg diluted in 0.9% saline, subcutaneous) 1 hour prior to anaesthesia with isoflurane (5% induction, ∼1% maintenance at 1 L/min with room air). Once anesthetised, a line block (bupivacaine, 0.2 ml of 0.25 mg/ml) was administered subcutaneously to the incision site. Saline (0.01 ml every 15 mins) was administered via intraperitoneal injection to maintain hydration throughout surgery. A midline sagittal incision was made to expose the skull before a high-speed drill was used to create bilateral craniotomies (0.8 mm diameter) overlying the right and left motor cortices (1.33 mm anterior and 2 mm lateral to Bregma – Allen Reference Atlas for adult C57BL/6J mice). Glass micropipettes were lowered into the craniotomies at depths of 250 μm and 400 μm from the brain surface to deliver the AAV into layers 2/3 and layer 5 respectively. A nanolitre injection system (World Precision Instruments Nanoliter 2020) was used to inject 200 nl of virus (≥7e12 vg/ml) at a rate of 40 nl/min, with the pipette left in place for 1 min before changing depth or removal. After all 4 injections were completed, the incision site was sutured closed (Surgical Specialties, Nylon 5-0 12 mm) and drops of topical anaesthetic (bupivacaine: 2.5 mg/ml) applied to the suture line. Mice were recovered on a heat pad and returned to their home cage for 24 days to allow for viral transfection.

### In vivo rTMS

For a more practical and translational approach to deliver chronic iTBS *in vivo*, the stimulation protocol for awake, freely moving mice differed from the *in vitro* stimulation. rTMS was delivered using the same rodent specific coils [30] used for the *in vitro* stimulation with the coil positioned to deliver peak stimulation over the primary motor cortices. The 600 pulses of the iTBS protocol [13] was delivered once a day for 7 consecutive days. The stimulation intensity generated a peak magnetic field of ∼0.2 T at the coil surface. Using electric field modelling of the exact same coil [4] estimates the stimulation parameters to induce a peak electric field of 30 V/m at the surface of the adult mouse brain [6].

As the mice were awake and freely moving during stimulation, were habituated to handling, the rTMS coil and noise from the power supply for 10 mins a day, 5 consecutive days prior to the first session of iTBS or sham. During handling, mice were accommodated to the placement of the coil on the surface of the head such that the windings of the coil were positioned over the primary motor cortices. Sham stimulation was identical to active iTBS, but the output of the waveform generator was not enabled. Stimulation was delivered by the same experimenter and at the same time each day (between 13:00 and 16:00).

### Euthanasia and tissue processing

Mice were deeply anesthetised with methoxyflurane (Penthrox) before lethal intraperitoneal injection of pentobarbital (0.15 ml of 160 mg/kg, Virbac) 3 hrs after the last stimulation session. After confirmation of death, mice were transcardially perfused with cold 0.9% saline followed by 2% paraformaldehyde (PFA) in 1X PBS for 10 mins at a flow rate of 10 ml/min. Brains were extracted and post-fixed in 2% PFA at 4 °C for 45 mins and stored at 4°C in 1X PBS containing 0.1% sodium azide. Brains were cryoprotected in sucrose (15% sucrose in 1X PBS for 24 hours followed by 30% sucrose in 1X PBS for a further 24 hours) and serially sectioned in the coronal plane at a thickness of 30 μm on a cryostat (Leica) for immunofluorescence.

### Brain slice staining and microscopy

Brain sections were washed with three 5 min washes in 1X PBS and blocked for 90 mins at room temperature in 1X PBS containing 0.1% Triton-X (100) and 10% normal goat serum. Sections were then washed with two 10 min 1X PBS washes and incubated for 2 hours at 4 °C in 1X PBS containing 0.1% Triton-X, 3% normal goat serum and Hoechst (1:1000, Invitrogen). Sections underwent three 5 min washes in 1X PBS and cover slipped with Prolong Diamond Antifade Mountant (Life Technologies). Coverslipped slides were stored at 4 °C for a minimum of 48 hours prior to imaging.

The same Nikon C2 confocal described above was used to image the brain sections using excitation and emission filters for DAPI (Hoescht), FITC (GFP), and TRITC (TdTomato). Two fields of view (60X magnification, 512 x 512 pixels) were captured for each cortical layer in brain sections that expressed TdTomato-labelled cell bodies. Multiple images were captured in the z-plane of each field of view, at a z-step depth of 0.25 um, capturing the appearance and disappearance of all AIS-GFP fluorescence in the field of view. Signal averaging of 2X was applied to image GFP to improve the signal-to-noise ratio. Images were pre-processed in FIJI and analysed in MATLAB as described for the *in vitro* analysis.

### Statistical analysis

A combination of Bayesian hierarchical linear models and estimation statistics were used to assess differences between groups for the *in vitro* and *in vivo* datasets. For estimation statistics, statistics and Cumming plots were generated on the raw data using resources described by Ho et al. [34].

Bayesian hierarchical linear models were done in R [35] using the brms package [36] and used log transformed AIS length and AIS distance data to improve normality of residuals, as evaluated using posterior predictive distributions and Q-Q plots. Each model included a continuous outcome variable (e.g., AIS length) predicted by a fixed-effects structure (Group for the *in vitro* experiments, and Group and Layer for *in vivo* experiments) as well as the interactions of any main effects. In addition, the models included random factors for coverslip nested in culture run (*in vitro*) and Animal_ID (*in vivo*). All models assumed Gaussian residuals, appropriate for the continuous nature of the outcome variables. Priors were placed on fixed effects (Normal (0, 5)) and on group-level standard deviations (Student t (3, 0, 2.5)). Models were fit using four Markov chains with 50,000 iterations each (10,000 warmup), for a total of 160,000 post-warmup samples. Convergence was assessed via effective sample size and Rhat diagnostics. Posterior estimates are reported as means with 90% credible intervals, defined as the range between the 5th and 95th percentiles of the posterior distribution.

Posterior samples were extracted and examined through pair plots to visualise parameter correlations and identify any divergent transitions. Model fit was further evaluated using posterior predictive checks to ensure the models adequately captured the data distribution. To aid interpretation, estimated marginal means (EMMs) were calculated for main effects and interactions using the emmeans package [37], with corresponding plots generated to visualise key findings. Bayesian hypothesis tests were performed using the hypothesis() function in brms to evaluate evidence for directional effects of predictors. For each effect, one-sided hypotheses were tested to quantify support for either an increase or decrease relative to the reference group. Bayes Factors (BF) derived from these tests were used to assess the strength of evidence for main effects and interactions.

BF are reported according to previously reported cut-offs that suggest evidence in favour of the alternative hypothesis:

- BF_10_ ≤1 = no evidence
- 1 < BF_10_ ≤ 3= anecdotal (weak) evidence
- 3 < BF_10_ ≤10= moderate evidence
- 10 < BF_10_ ≤100= strong evidence
- BF_10_ >100= extreme evidence

To aid in the interpretation of our analysis, we set a BF_10_ threshold of ≥3 to suggest a meaningful effect [38].

## Results

### 6 hours of iTBS or KCl stimulation *in vitro* causes small shifts to AIS distance from the soma

To provide proof of principle that chronic iTBS could promote structural AIS plasticity in cortical neurons, we used a primary neuronal cell culture model, immunolabelling for Ankyrin G to identify the AIS, MAP2 to identify the soma and dendrites, GAD65/67 to identify inhibitory neurons, and Hoechst dye to identify the nucleus (Fig 1). Given that the effects of rTMS often extend beyond the period of stimulation, we investigated structural changes to the AIS immediately post-stimulation (+0hrs), and in a second group where the cultures remained in the incubator for 24 hours post-stimulation (+24hrs).

**Figure 1.**
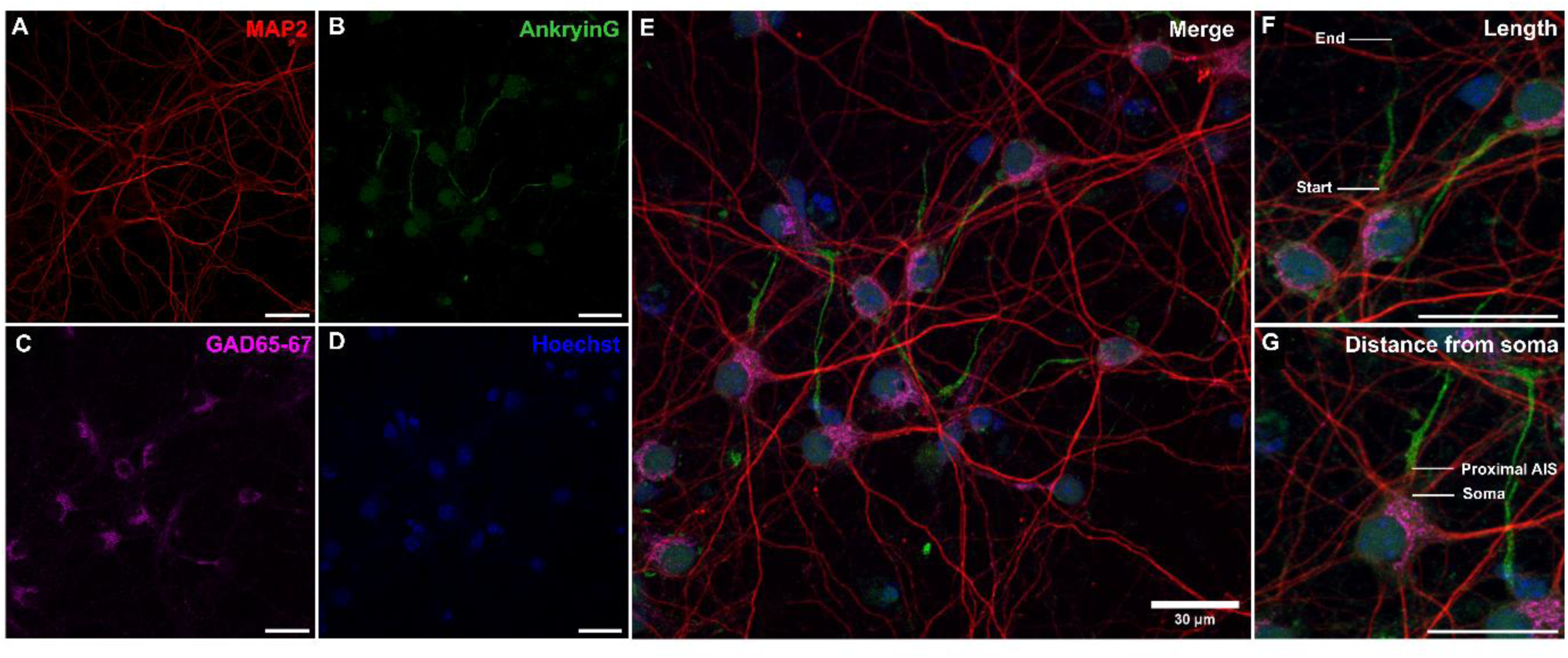
Maximum intensity z-projection of immunolabelled cortical inhibitory neurons at DIV7. Cortical neurons were immunolabelled for the neuronal marker MAP2 (A), the AIS marker Ankyrin G (B), the inhibitory marker GAD65-67 (C), and the nuclear marker Hoechst (D). Z-projection images were merged (E) before measurements of AIS length (F) and AIS distance from soma (G) were taken. Images were captured on a confocal microscope at 60x magnification. All scale bars =30 µm.

At + 0hrs, rMS caused a 1.34 µm distal shift in AIS position from the soma relative to sham-stimulated neurons (BF = 232; sham: 6.26 µm [95% CI: 5.73 µm – 6.84 µm]; rMS: 7.60 µm [95% CI: 6.73 µm – 8.59 µm]; Fig 2 A-B), which reversed to a proximal shift of 1.22 µm at + 24 hrs (BF = 61.0; sham: 6.80 µm [95% CI: 6.18 µm – 7.49 µm]; rMS: 5.58 µm [95% CI: 4.94 µm – 6.32 µm]; Fig 2 A, C) relative to sham. For KCl, there was no change to AIS distance from soma at both +0hrs (BF= 2.82; KCl= 5.90 µm [95% CI: 5.35 µm – 6.51 µm]) or +24 hrs post stimulation (BF = 1.2; sham: 7.05µm [95% CI: 6.38 µm – 7.79 µm], KCl: 7.53 µm [95% CI: 5.75 µm – 9.89 µm]; Fig 2 A, C). Together, these findings suggest that 6 hours of subthreshold iTBS but not 15 mM KCl altered AIS distance relative to the soma in primary cortical inhibitory neurons that was dependent on the time post-stimulation.

**Figure 2.**
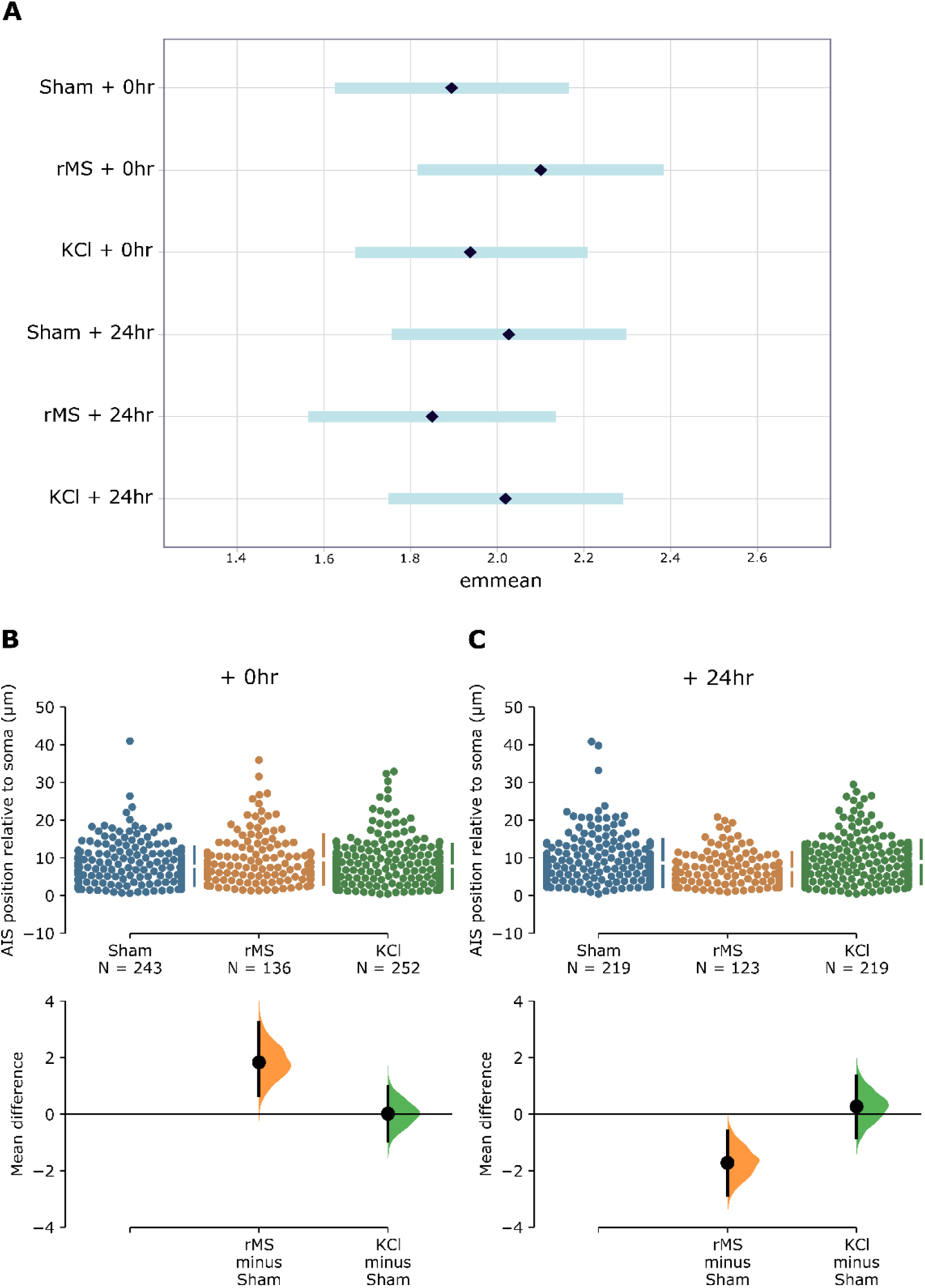
Model-adjusted means (EMMeans) and raw data (Cumming estimation plots) of AIS position relative to the soma following 6 hours of rMS or KCl stimulation. EMMeans represent model-adjusted group effects, while estimation plots display individual data points and group mean differences. (A) EMMeans of AIS position relative to the soma plotted with 95% credible intervals. (B) Cumming estimation plots of raw data showing individual data points (top) and mean group differences (bottom) plotted with 95% confidence intervals for +0hrs and (C) +24hrs. Number of cells indicated by “N” with the data at each timepoint collected from at least 12 coverslips over a minimum of 6 separate culture runs. Each culture run contained neurons collected from a minimum of 3 P1 mice.

### 6 hours of iTBS or KCl stimulation *in vitro* induces structural plasticity of AIS length

We next investigated whether 6 hours of rMS or KCl stimulation altered AIS length. Our analysis showed that rMS had no effect on AIS length at +0hrs (BF = 2.12; sham: 24.53 µm [95% CI: 22.90µm - 26.30µm]; rMS: 24.86 µm [95%CI: 23.06 µm – 26.82 µm], Fig 3 A-B). However, at +24hrs, rMS AIS lengths were shorter by 4.05 µm relative to sham stimulated neurons (BF = ∞; sham: 26.57 µm [95 % CI: 24.82 µm – 28.49 µm]; rMS: 22.52 µm [95% CI: 20.86 µm – 24.29 µm]; Fig 3 A, C).

**Figure 3.**
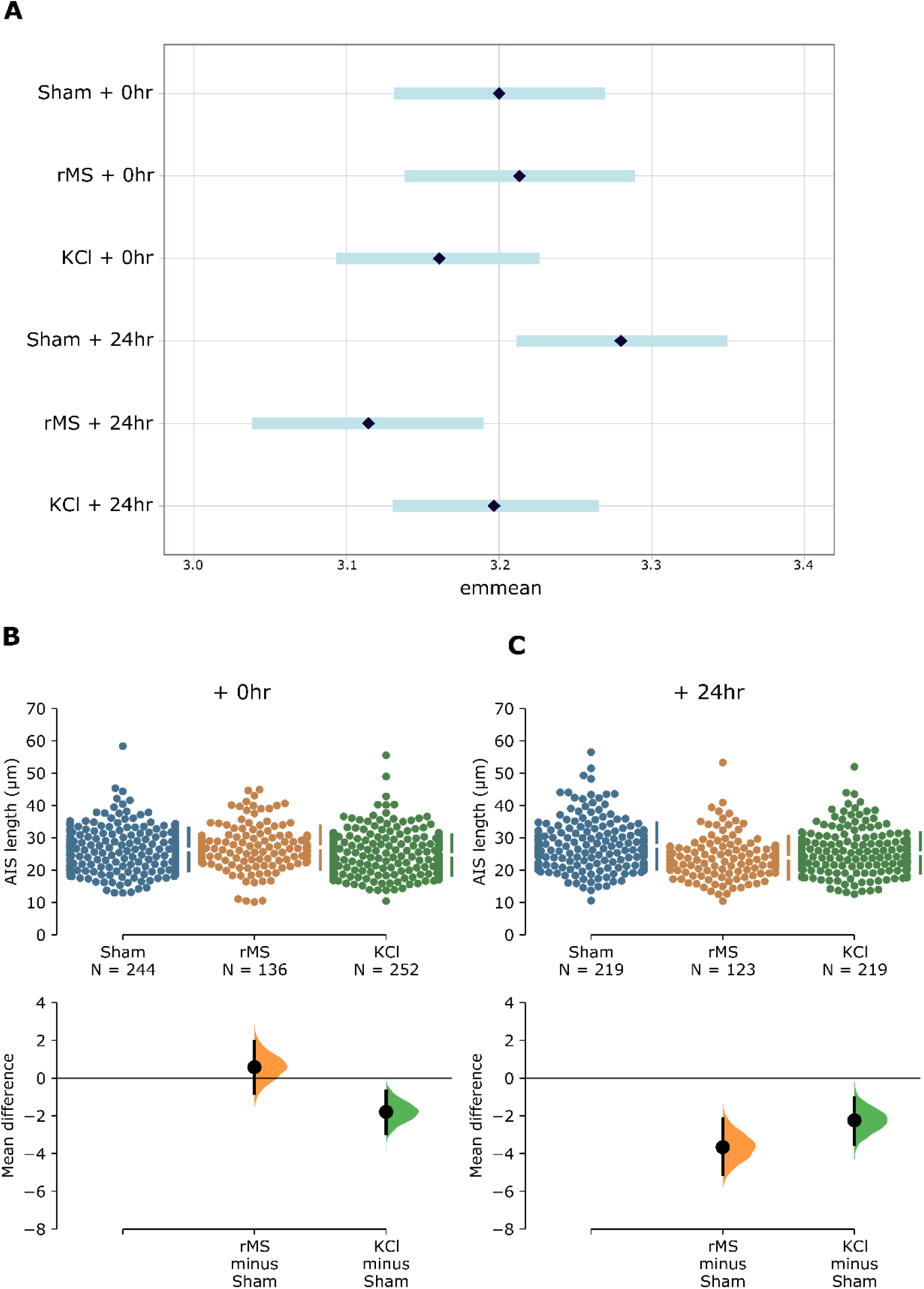
Model-adjusted means (± 95% CI; Emmeans) and raw data (Cumming estimation plots) of AIS length following 6 hours of rMS or KCl stimulation. Emmeans represent model-adjusted group effects, while estimation plots display individual data points and group mean differences. (A) Emmeans of AIS length plotted with 95% credible intervals. (B Cumming estimation plots of raw data showing individual data points (top) and mean group differences (bottom) plotted with 95% confidence intervals for +0hrs and (C) +24hrs. Number of cells indicated by “N” with the data at each timepoint collected from at least 12 coverslips over a minimum of 6 separate culture runs. Each culture run contained neurons collected from a minimum of 3 P1 mice.

In contrast, AIS lengths were shorter relative to sham at both time points for KCl stimulated neurons, with a 0.94 µm reduction at +0hrs (BF = 15.4; KCl: 23.59 µm [95% CI: 22.05 µm – 25.19 µm; Fig 3 A-B) and a 2.12 µm reduction at +24hrs (BF = 2856; KCl: 24.45 µm [95% CI: 22.87 µm – 26.19 µm]; Fig 3 A, C). These findings suggest that 6 hours of rMS induces delayed shortening of AIS lengths in primary cortical inhibitory neurons whereas KCl stimulation induces acute and long-lasting shortening of AIS lengths.

### Molecular mechanisms of rMS-induced structural AIS plasticity

To identify the molecular mechanisms underlying the structural AIS plasticity changes we observed in primary cortical neurons, we blocked L- and T type Ca_v_ channels with mibefradil, or Na_v_ channels with TTX during sham, rMS, and KCl stimulation in a separate group of coverslips.

In the presence of mibefradil, rMS continued to induce a distal shift in AIS position (+2.21 µm) from the soma relative to sham at +0hrs (BF = 29.9; sham: 7.72µm [95% CI: 6.69µm – 8.90µm]; rMS: 9.93µm [95% CI: 8.40µm – 11.7µm]; Fig 4 A-B). Interestingly, the reversal to a proximal shift at +24hrs was prevented in the presence of mibefradil (BF = 0.42; sham: 7.85µm [95% CI: 6.93µm – 8.88µm]; rMS: 7.70µm [95% CI: 6.64µm – 8.93µm]; Fig 4 A, C).

**Figure 4.**
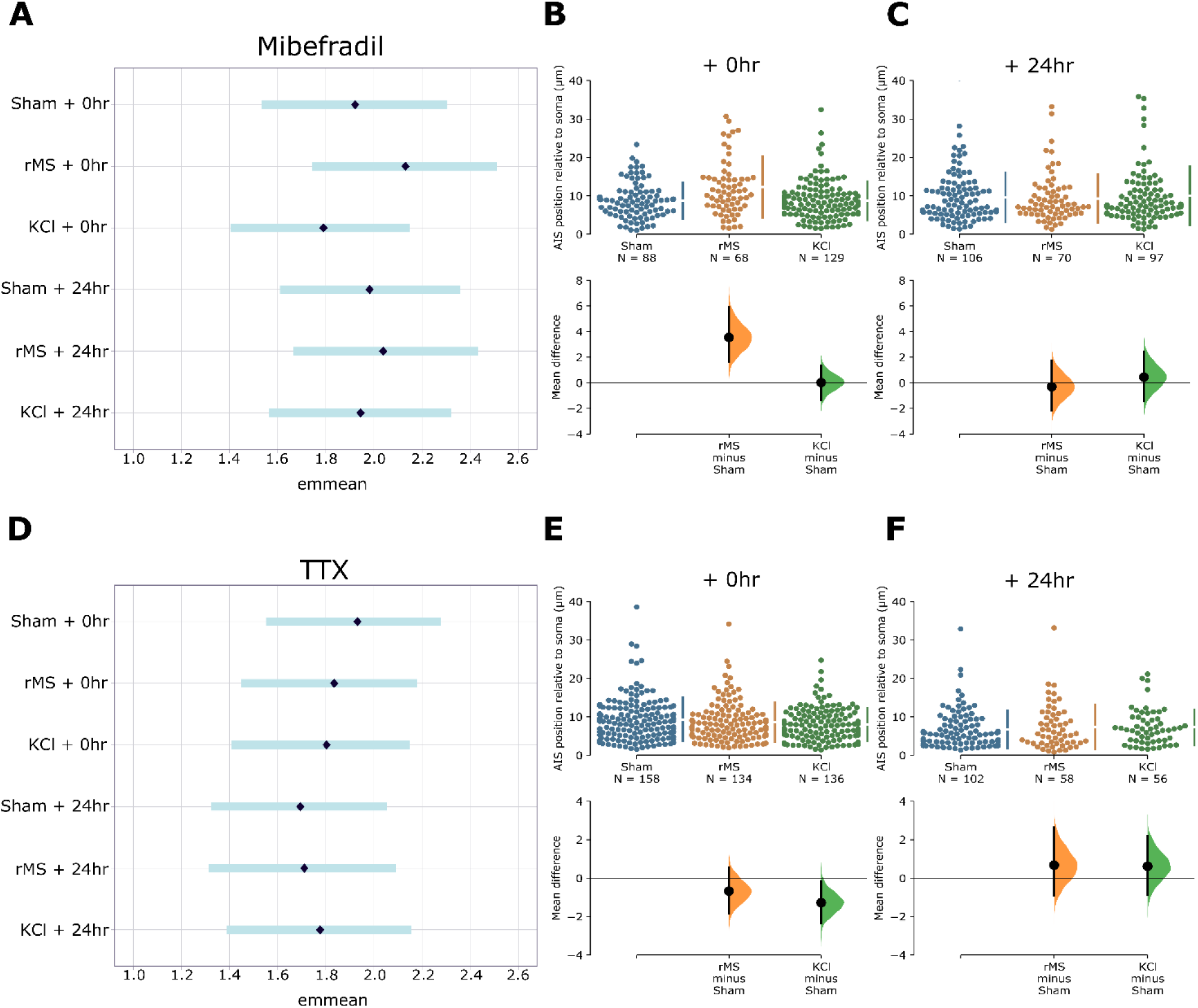
Model-adjusted means (± 95% CI; Emmeans) and raw data (Cumming estimation plots) of AIS position relative to the soma following 6 hours of rMS or KCl stimulation in the presence of mibefradil or TTX. Emmeans represent model-adjusted group effects, while estimation plots display individual data points and group mean differences. (A) Emmeans plotted with 95% credible intervals of AIS distance from soma when incubated with mibefradil. (B) Raw data of AIS distance relative to the soma measurements collected from individual neurons (top) and mean differences of treatment group compared to sham group (bottom) plotted as bootstrap sampling distributions with 95% confidence intervals at + 0hrs and (C) +24 hrs. (D) Emmeans plotted with 95% credible intervals of AIS distance from soma when incubated with TTX. (E) Raw data of AIS position relative to the soma and mean differences of treatment groups compared to sham in the presence of TTX at + 0 hrs and (F) +24 hrs. Number of cells is represented by “N” with data from each timepoint collected from at least 6 coverslips over a minimum of 3 separate culture runs. Each culture run contained neurons collected from a minimum of 3 P1 mice.

Blocking with TTX prevented the distal shift in AIS position by rMS at +0hrs, leading to a very small proximal shift in AIS position (0.47µm) from the soma relative to sham (BF = 7.69; sham: 7.82µm [(95% CI: 7.11µm – 8.61µm]; rMS: 7.35µm [95% CI: 6.69µm – 8.09µm]; Fig 4 D-E), and proximal shift in AIS position at +24hrs (BF = 1.24; sham: 5.78µm [95% CI: 4.81µm – 6.96µm]; rMS: 5.49µm [95% CI: 4.50µm – 6.70µm]; Fig 4 D, F).

From these experiments we show that the acute distal shift in AIS position induced by rMS is Na_v_ channel dependent, whereas the delayed proximal shift in AIS position requires both Na_v_ and L and T-type Ca_v_ channels. This contrasts with KCl where the acute distal shift in AIS position was Na_v_ and L and T-type Ca_v_ channel dependent.

For changes to AIS length, incubation with mibefradil during the 6 hours of rMS caused a 2.27µm increase in AIS lengths compared to sham that was not present with rMS alone (BF = 27.8; sham 24.98 [95% CI: 22.22µm – 28.08µm]; rMS 27.25µm [95% CI: 24.10µm – 30.61µm]; Fig 5 A, C). In the rMS +24hrs group, mibefradil did not completely prevent the shortening of AIS length as AIS lengths were 2.15 µm shorter compared to sham (BF = 20.0; sham 27.83µm [95% CI: 24.80µm – 31.27µm]; rMS 25.68µm [95% CI: 22.70µm – 28.94µm]; Fig 5 A, C).

**Figure 5.**
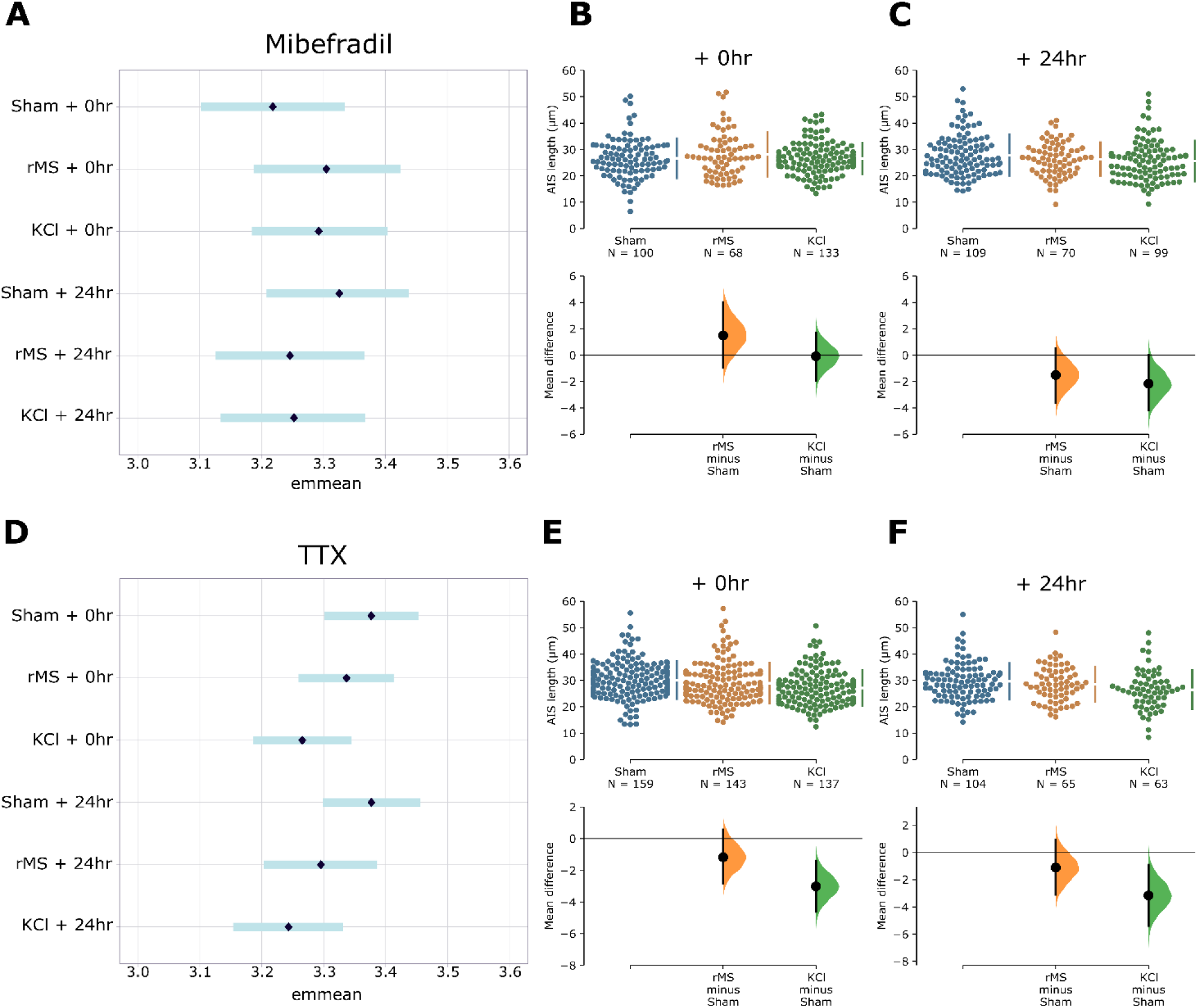
Model-adjusted means (± 95% CI; emmeans) and raw data (Cumming estimation plots) of AIS Length following 6 hours of rMS or KCl stimulation in the presence of mibefradil or TTX. Emmeans represent model-adjusted group effects, while estimation plots display individual data points and group mean differences. (A) Emmeans plotted with 95% credible intervals of AIS lengths when incubated with mibefradil. (B) Raw data of AIS length measurements collected from individual neurons (top) and mean differences of treatment group compared to sham group (bottom) plotted as bootstrap sampling distributions at + 0 hours and (C) + 24 hours post stimulation. (D) Emmeans plotted with 95% credible intervals of AIS lengths when incubated with TTX. (E) Raw data of AIS lengths and mean differences of treatment groups compared to sham in the presence of TTX at + 0 hrs and (F) +24 hours post stimulation. The total number of neurons analysed from each group is presented under the group names. Data from each treatment group at each timepoint was collected from at least 6 coverslips over a minimum of 3 separate culture runs. Each culture run contained neurons collected from a minimum of 3 P1 mice.

Incubating mibefradil with KCl prevented AIS shortening at +0hrs with AIS lengths 1.93µm longer than sham stimulated neurons (BF = 36.4; KCl: 26.91µm [95% CI: 24.10 – 30.02µm]; Fig 5 A-B). However, blocking with mibefradil did not prevent the KCl induced shortening of AIS lengths at +24hrs, with AIS lengths 1.98 µm shorter than sham stimulated neurons (BF = 38.2; KCl: 25.85µm [95% CI: 23.00µm – 29.08µm]; Fig 5 A, C).

Unexpectedly, rMS coverslips incubated with TTX caused a 1.85 µm shortening in AIS length at +0hrs that was not present with rMS alone (BF=10.40, sham: 29.28 µm [95% CI: 27.14µm – 34.60µm]; rMS 28.13µm [95% CI: 26.04µm – 30.38µm]; Fig 5 D, E) with subsequent analysis comparing sham without TTX to sham with TTX showing that the addition of TTX alone caused AIS plasticity. Nevertheless, incubation with TTX partly blocked rMS induced shortening of the AIS at +24hrs relative to sham, reducing the effect to 2.29µm (BF=38.4, sham: 29.28 µm [95% CI: 27.06µm – 31.10µm]; rMS 26.99µm [95% CI: 24.60µm – 29.55µm]; Fig 5 D, F).

For KCl stimulated coverslips, blocking with TTX did not prevent KCl induced AIS shortening relative to sham stimulated neurons at either timepoint, with the effect size increasing at +0hrs to a 3.09 µm shortening (BF = 6666; KCl: 26.19µm [95% CI: 24.20µm –28.36 µm]; Fig 5 D-E) and a 3.66 µm shortening at +24hrs (BF = 2024; KCl: 25.62µm [95% CI: 23.42µm – 27.98µm]; Fig 5 D, F).

From these experiments, we find that delayed AIS length shortening with rMS is partially dependent on Na_v_ and L and T Ca_v_ channels. Whereas KCl induced AIS length shortening, is not Na_v_ channel dependent and only dependent on L- and T-type Ca_v_ channels for the acute change.

### Chronic rMS or KCl stimulation *in vitro* does not promote neuronal injury

As neuronal injury can cause maladaptive AIS structural plasticity [39,40], a separate group of coverslips were used to assess cell viability following chronic stimulation to quantify the number of cells with compromised cell membranes. Our analysis showed that chronic rMS did not increase the percentage of damaged/dying neurons at +0hrs (BF=0.10, sham: 11.49% [95% CI: 3.68% – 33.15%]; iTBS: 3.79% [95% CI: 0.22% – 17.30%]; Fig 7 A-B) or +24hrs (BF=1.09, sham: 7.05% [95% CI: 1.10% – 30.18%]; iTBS: 7.51% [95% CI: 1.18% – 31.77%]; Fig 7 A-C). This was similar for KCl, with no increase to the percentage of damaged/dying neurons relative to sham at +0hrs (BF=0.86,KCl: 10.93% [95% CI: 2.93% – 34.12%]; Fig 7 A-B) or +24hrs (BF=1.14, KCl: 7.61% [95% CI: 0.93% – 41.10%]; Fig 7 A-C).

**Figure 6.**
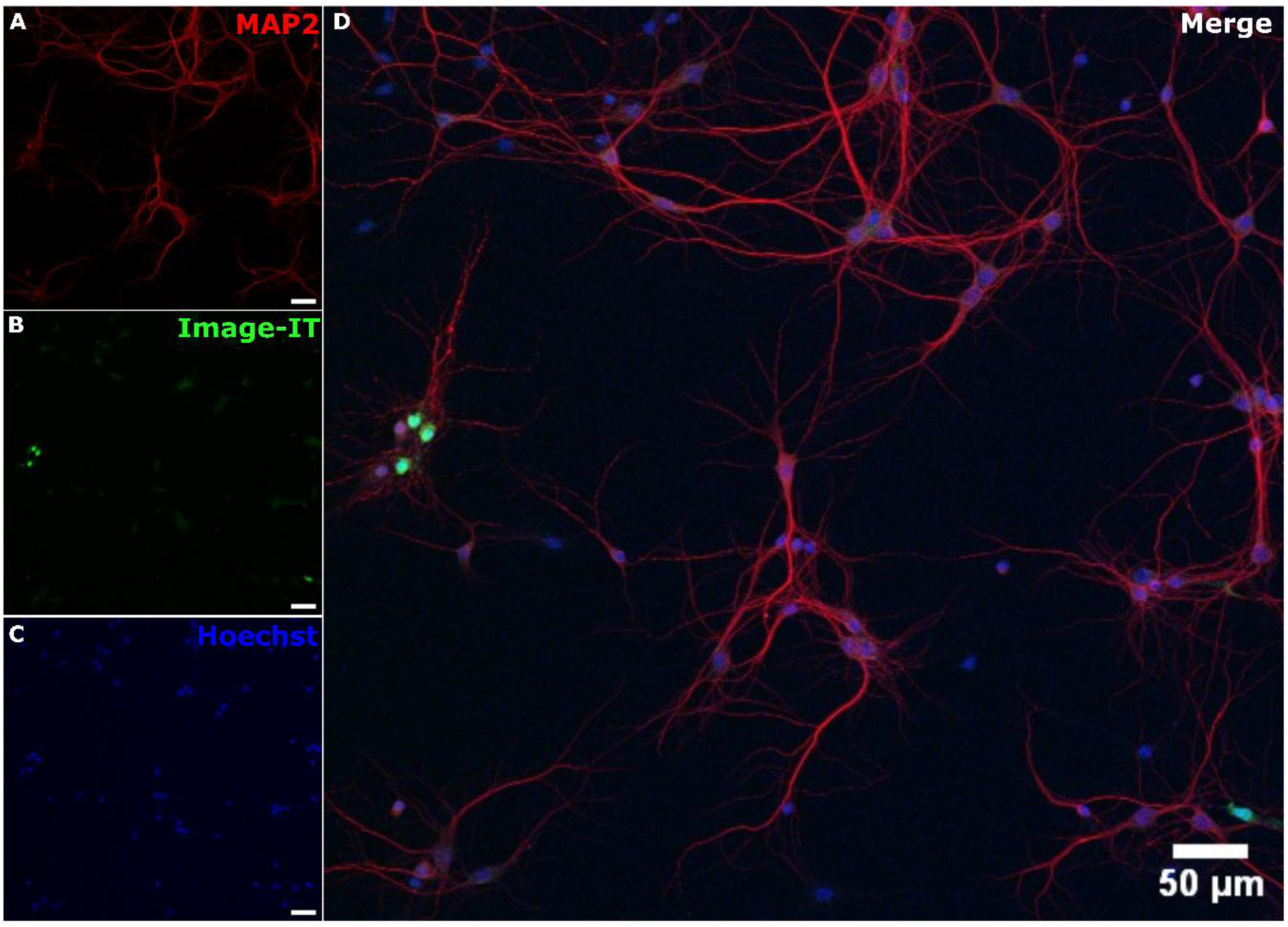
Representative image of the Image-IT assay. (A) Immunolabelling of MAP2 to label the soma and dendrites in red. (B) Cells stained with the green Image-IT dye identifying cells with compromised cell membranes. (C) Hoechst stain labelling identifying cell nuclei. (D) Merge of MAP2, Image-IT, and Hoechst labelling to identify dying/damaged cells from non-dying/damaged cells. All scale bars= 50µm.

**Figure 7.**
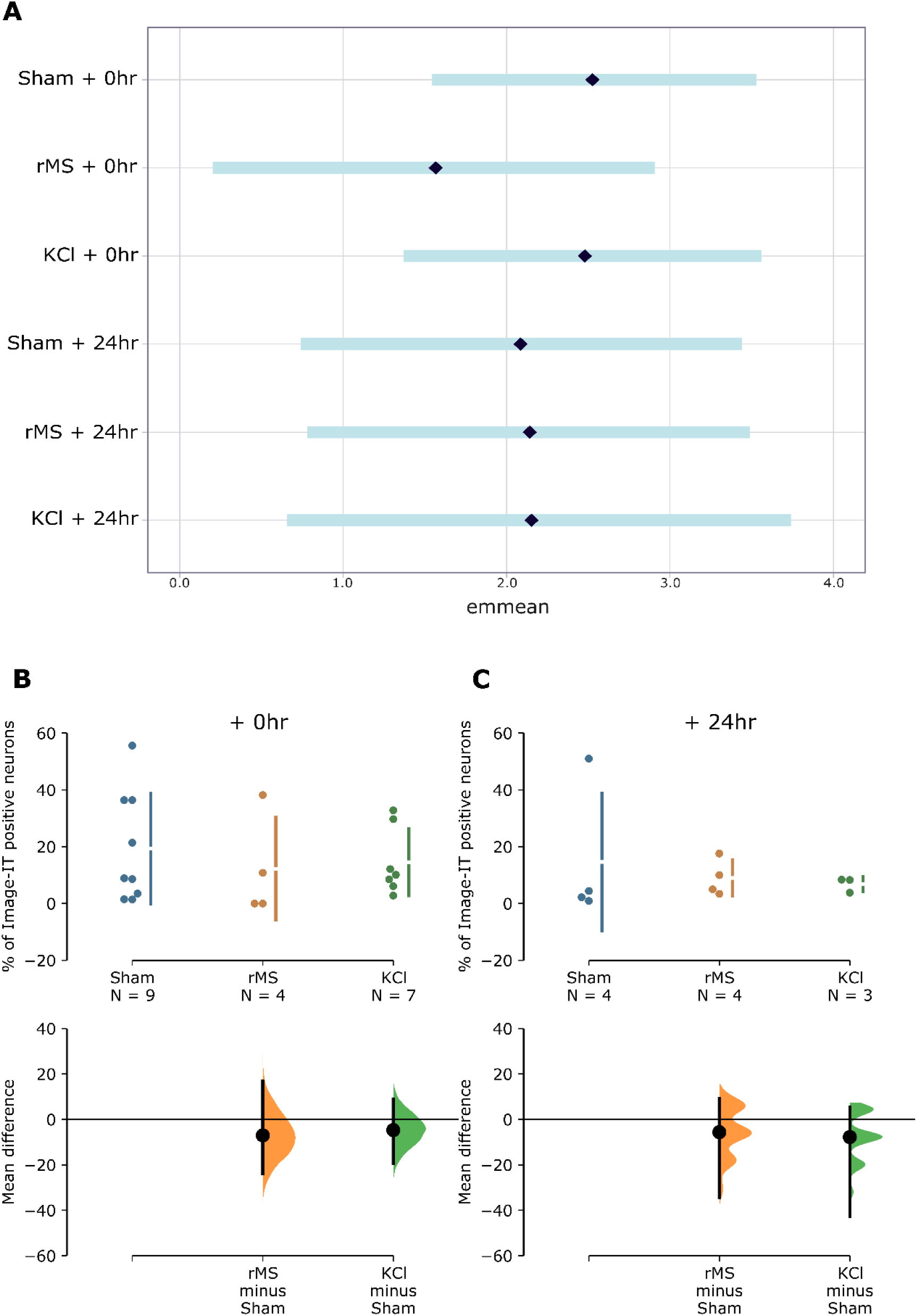
Model-adjusted means (± 95% CI; emmeans) and raw data (Cumming estimation plots) of percent of Image-IT positive neurons following 6 hours of rMS or KCl stimulation. Emmeans represent model-adjusted group effects, while estimation plots display individual data points and group mean differences. (A) Emmeans and 95% crddible intervals of transformed percentage of Image-IT positive/ cells with compromised membranes following rMS or KCl relative to sham at +0hrs or +24hrs. (B) Raw data of Image-IT collected at +0hrs and (C) +24hrs presented as number of positive cells in each coverslip (top) and mean differences of treatment group compared to sham group (bottom) plotted as bootstrap sampling distributions. N represents the number of coverslips per group with the data from each treatment group and timepoint collected from at least 1 coverslip over a minimum of 3 separate culture runs. Each culture run contained neurons collected from a minimum of 3 P1 mice.

These findings suggest that the AIS structural plasticity induced by chronic rMS and KCl *in vitro* is not due to neuronal injury.

### Daily iTBS for 7 consecutive days reduces AIS length in cortical neurons of the motor cortex

Having established a proof of principle that chronic iTBS could induce AIS plasticity in cortical neurons *in vitro*, we next investigated whether this could be translated *in vivo* by adapting to a more translational paradigm of daily iTBS (600 pulses) for 7 consecutive days. Analysis of AIS lengths in cortical neurons (Fig 8) in layers 2/3 and 5 showed a modest reduction of 1.35µm relative to sham (BF=6.08, iTBS: 23.55µm [95% CI: 21.54µm – 25.74µm]; sham: 24.90µm [95% CI: 22.77 µm – 27.31µm]; Fig 9 A-B). This effect was consistent across cortical layers 2/3 (sham: 25.67 µm; iTBS: 23.88 µm) and 5 (sham: 24.18 µm; iTBS: 23.20 µm) (Fig 9 C-E).

**Figure 8.**
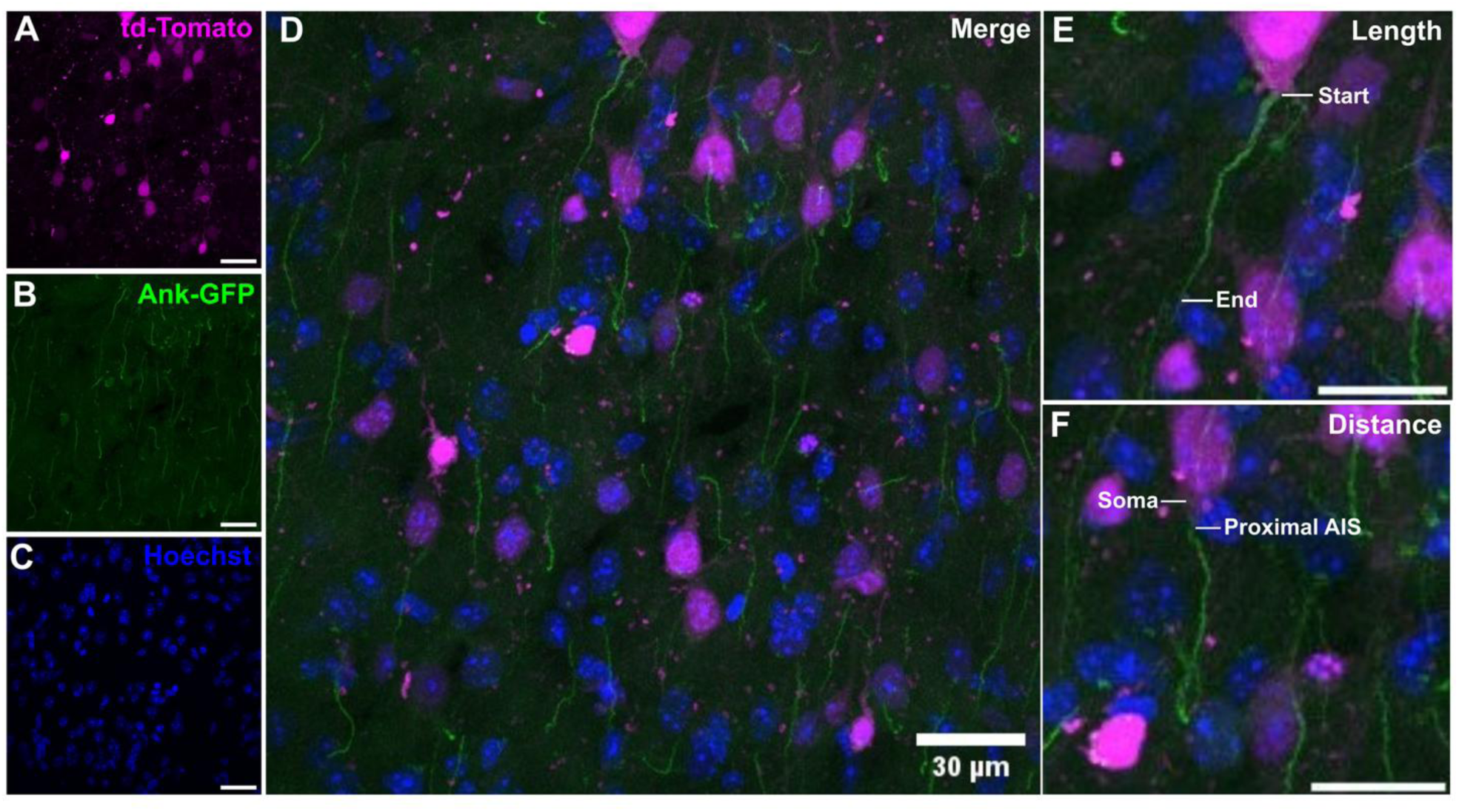
Maximum intensity z-projections of confocal microscopy capturing layer 5 pyramidal neurons of the mouse motor cortex. AnkG-GFP mice were injected AAV-hSynapsin-Cre-TdTomato to induce GFP expression at the AIS of cortical pyramidal neurons in layers 2/3 and layer 5 of the motor cortex. (A) td-Tomato labelled neuronal cell bodies and dendrites, (B) GFP labelling, and (C) Hoechst labelled cell nuclei. (D) Merged image of the three channels to show (E) AIS length (F) and distance from the cell body. All images acquired at 60x magnification and scale bars = 30µm.

**Figure 9.**
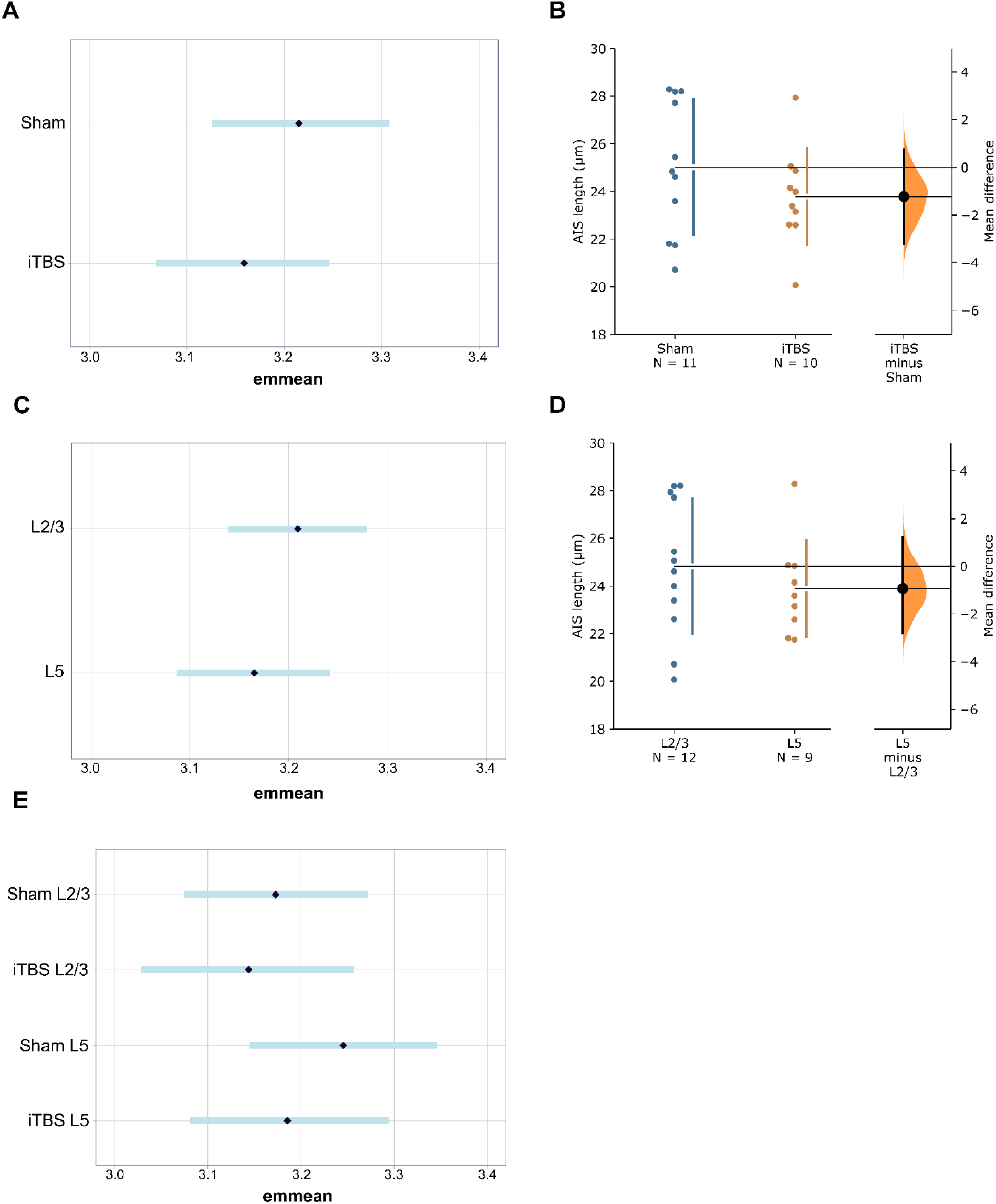
Model-adjusted means (±95% CI; emmeans) and raw data (Cumming estimation plots) of AIS length. Emmeans represent model-adjusted group effects, while estimation plots display individual data points and group mean differences. (A) Emmeans plotted with 95% credible intervals of AIS lengths irrespective of cortical layer. (B) Raw data of AIS lengths presented with each dot representing the average AIS length in each mouse analysed plotted with 95% bootstrap confidence intervals. (C) Emmeans plotted with 95% credible intervals of AIS lengths in cortical layers 2/3 and 5 irrespective of stimulation condition. (D) Raw data of AIS lengths of the average AIS lengths in cortical layers 2/3 and 5 from each mouse analysed plotted with 95% bootstrap confidence intervals. (E) Emmeans plotted with 95% credible intervals of AIS lengths separated by stimulation condition and cortical layer. N represent the number of mice with a minimum of 120 AIS lengths quantified per mouse.

From our images, we were only able to accurately measure distance from Td-Tomato positive cell bodies in a relatively small number of cells across all groups, preventing informative statistical analysis. Based on the EMmeans alone, neurons from iTBS mice had a more proximal AIS (−1.58um: see Table 1 for summary).

**Table 1.**
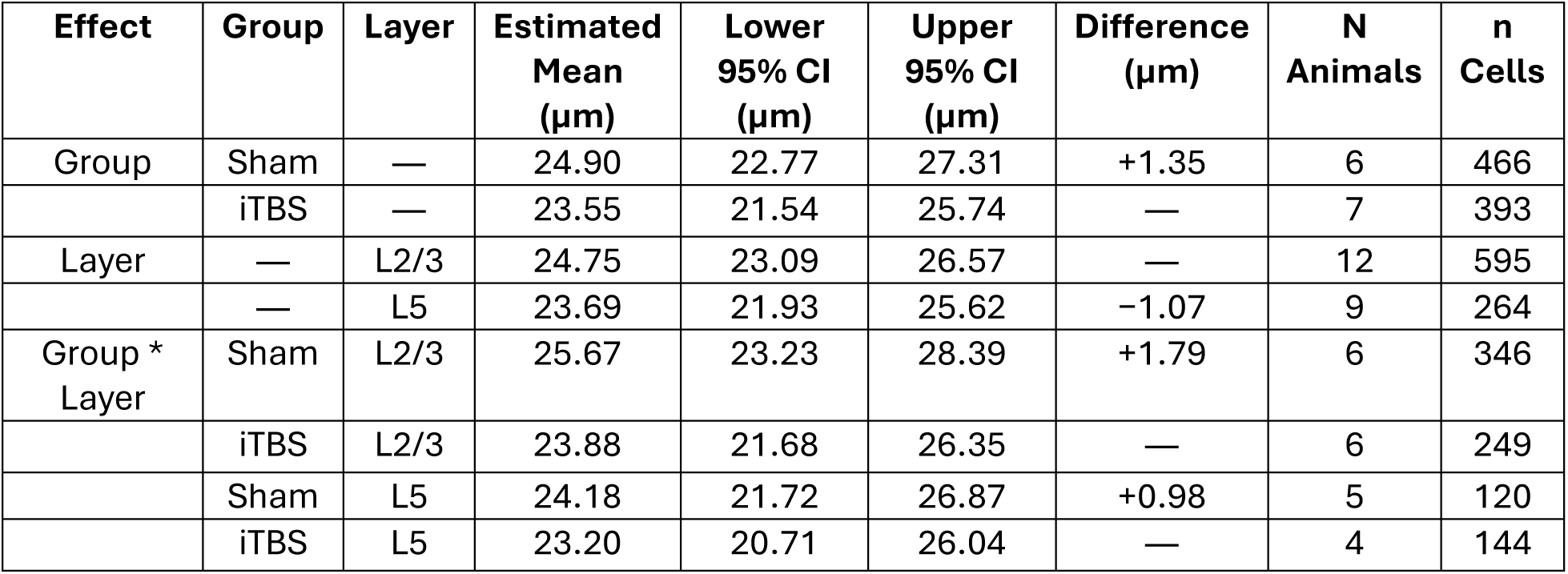
*In vivo* quantification of AIS distance from soma.

## Discussion

Using primary cell cultures and an *in vivo* mouse model, we show that chronic stimulation with iTBS induces structural AIS plasticity in cortical neurons. As hypothesised, chronic iTBS induced structural AIS plasticity in a direction that reduces neural excitability. In cultured neurons, 6 hours of iTBS caused an initial Na_v_ channel dependent distal shift in AIS position relative to the soma that reversed to a Na_v_ and L and T-type Ca_v_ channel dependent proximal shift 24 hours post-stimulation. The changes to AIS position *in vitro* were accompanied by a shortening of AIS length 24 hours post-stimulation that was partly L and T-type Ca_v_ channel dependent. Critically, the structural AIS plasticity induced *in vitro* was not due to neuronal injury, as cell viability assays confirmed that the 6-hour protocol of iTBS used did not increase the amount of dying or damaged cells relative to sham. Moving to an *in vivo* model, we found that daily iTBS for 7 consecutive days to the primary motor cortex shortened the AIS length in layer 2/3 and 5 pyramidal neurons.

Compared to other non-invasive brain stimulation techniques, we find differences to the effect on AIS structural plasticity and their effect sizes. In cultured inhibitory neurons at the same DIV, 6 hours of static magnetic stimulation reduced AIS lengths at +0hrs and +24hrs by ∼2µm that was not dependent on L and T-type Ca_v_ channels, with no changes to AIS location [28]. In contrast, we found that 6 hours of iTBS reduced AIS lengths by 3.6µm at +24hrs that was partly dependent L- and T-type Ca_v_ channels and caused a bidirectional and time-dependent change to AIS location suggesting that repetitive magnetic is more suitable for inducing changes to AIS location and larger changes to AIS length than static magnetic stimulation. Compared to optogenetic stimulation of hippocampal neurons *in vitro* [32], chronic subthreshold iTBS reduced the AIS length to a similar extent (3.5 µm with optogenetics), but it should be noted that AIS plasticity may vary between cell types (discussed further below).

For our *in vivo* experiments, we adapted the iTBS protocol used *in vitro* to a more translational paradigm that could be used in clinical and non-clinical populations. While the AIS shortening induced with daily iTBS was small (−1.4µm), this structural change is still likely to lead to a functional change as combined patch clamp electrophysiology and live imaging of CA1 pyramidal neurons from AnkG-GFP mice suggests a linear relationship between changes in action potential threshold and AIS length, with a ∼0.5mV change in action potential threshold for every 1µm change in AIS length [27]. Extrapolating this relationship to our cortical pyramidal neurons, also from AnkG-GFP mice, we estimate a 0.7mV increase in action potential threshold. While modest, this decrease in neural excitability within the motor cortex could have significant effects at the network level, especially as the reduced AIS length occurred in the main input and output layers of the motor cortex. Interestingly, the reduction in AIS length following iTBS *in vivo* was similar to the 1.5µm reduction to dentate granule cells AIS lengths following 2 hours of suprathreshold electrical stimulation in anaesthetised rats [25]. In that study, computational modelling suggested the small change in AIS length would increase action potential latency and amplitude, decreasing intrinsic excitability. Taken together, the results suggest 7 consecutive days of daily iTBS would decrease intrinsic excitability, and that greater stimulation intensity alone in brain stimulation does not guarantee larger effect sizes.

Similar to previous studies [15,32,41,28,23], we used KCl stimulation as a positive control to induce structural AIS plasticity. In cultured cortical inhibitory neurons, we found that 15mM KCl stimulation for 6 hours did not alter AIS position relative to the soma, but shortened AIS length at +0 and +24hrs post-stimulation. This replicated the findings from our previous paper investigating the effect of 6 hours of 15mM KCl or static magnetic stimulation on AIS structural plasticity in cultured cortical inhibitory neurons [28]. Together with the present study, we suggest that 15mM KCl stimulation is a useful experimental tool to study changes to AIS length in cortical inhibitory neurons, but not for changes to AIS position. However, KCl induced AIS structural plasticity is likely to be cell type specific. For example, the seminal *in* vitro AIS plasticity paper by Grubb and Burrone [15] showed 48 hours of 15mM KCl stimulation induced a distal AIS shift in excitatory but not inhibitory hippocampal neurons. Similarly, 3 hours of 15mM KCl stimulation to cultured hippocampal dentate granule neurons shortened AIS lengths in CA3 but not CA1 neurons [32]. Therefore, our findings further highlight the need to consider cell type when using KCl as a positive control.

A surprising finding from experiments was the induction of AIS structural plasticity with the addition of mibefradil and TTX in some groups. The addition of TTX reduced AIS lengths in the sham stimulated coverslips, whereas mibefradil increased AIS lengths in sham stimulated coverslips. To our knowledge, neither TTX or mibefradil has been shown to induce AIS plasticity previously and we can only speculate on the mechanism for such changes. In the case of TTX, silencing spontaneous neural activity has been shown to promote compensatory synaptic mechanisms to increase neural activity, with increased neurotransmitter release [42] and phosphorylation of proteins involved in synaptic processes [43]. Therefore, the reduction in AIS length following TTX may be a homeostatic mechanism to stabilise neural activity following prolonged synaptic enhancement. In contrast, we speculate mibefradil reduces spontaneous neural activity [44] by blocking calcium entry from subthreshold membrane depolarisations, causing a homeostatic increase in AIS length to increase overall activity. However, irrespective of the mechanism, our findings should be considered by future studies using TTX or mibefradil to dissect AIS plasticity mechanisms, particularly for inhibitory cortical neurons.

A limitation of our experiments was the inability to quantify AIS distance from the soma in our virus labelled tissue. This was due to an inability to confidently identify the edge of the soma in a large number of cells for all animals. However, it is worth noting that changes to AIS position are rarely seen *in vivo*, with behavioural [17], pharmacological [23,27], and brain stimulation interventions [25] observing changes to AIS length but not distance. Therefore, even with larger sample sizes per animal, it is unlikely that a change to AIS position would have been observed. Lastly, we acknowledge that the repetitive magnetic stimulation protocols we used *in vitro* and *in vivo* may not be the optimal protocols for inducing AIS plasticity. Further studies are needed to determine whether the direction and effect size of AIS plasticity induced *in vivo* vary between stimulation frequencies, amount of stimulation (e.g., accelerated rTMS over days vs single daily sessions rTMS over weeks), brain region, and age. This is particularly important given that recent studies using spatial transcriptomics have shown that the neural plasticity mechanisms induced by rTMS differ between rTMS protocol, age, and brain region, down to individual cortical layers [6,9]

In conclusion, we provide further evidence that repetitive magnetic stimulation can induce neural plasticity beyond the synapse, with changes to structural AIS plasticity, an effect known to underpin intrinsic plasticity. These findings expand the use of rTMS to study AIS structural plasticity non-invasively and potentially as a treatment for neurological disorders involving abnormal AIS structural plasticity such as motor neurone disease [45] or dementia [46,47]. Furthermore, our findings highlight the need to consider the recruitment of non-synaptic homeostatic neural plasticity mechanisms when delivering rTMS for multiple days or weeks.

## Acknowledgements

The authors thank Professor Paul Jenkins (University of Michigan) for gifting the initial AnkG-GFP mice used to establish a colony at the University of Western Australia for axon initial segment research. This project was funded by a Sarich Family of Western Australia Fellowship to ADT. EKS was supported by a Research Training Program scholarship from the Australian government, and a Byron Kakulas top-up scholarship from the Perron Institute for Neurological and Translational Sciences.

## Author contributions

EKS: Data curation, formal analysis, investigation, visualisation, writing – review & editing

LJA: Investigation

WCA: Writing – review & editing

JR: Funding acquisition, writing – review & editing

JNJR: Writing – review & editing

DC: Conceptualisation, investigation, methodology, writing – review & editing, supervision

JLB: Conceptualisation, Investigation, methodology, project administration, supervision, writing – review & editing

ADT: Conceptualisation, formal analysis, investigation, visualisation methodology, project administration, funding acquisition, supervision, writing – original draft, writing – review & editing

