## Supplementary material for "Chronic subthreshold intermittent theta burst stimulation promotes structural axon initial segment plasticity in cortical neurons"

### Supplementary Materials

Figure S.1: Approximation of induced electric field distribution across the neuronal layer of the coverslip during each iTBS pulse.

Table S.1 Categorisation of neuronal subtypes in the cell cultures.

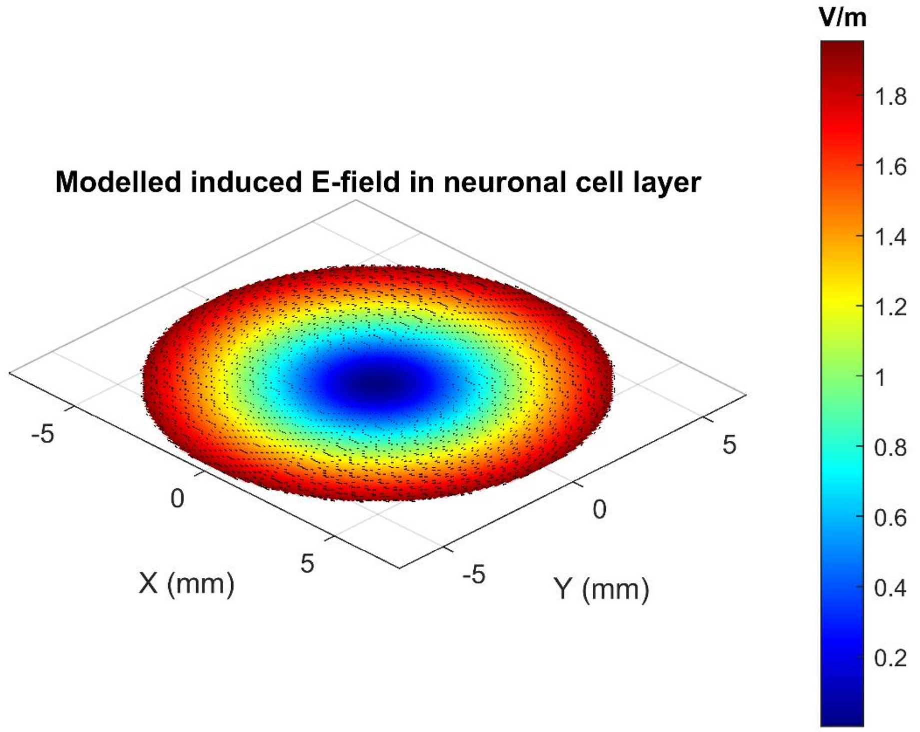


**Figure S.1: Approximation of induced electric field distribution across the neuronal layer of the coverslip during each iTBS pulse.** Peak electric field intensity of 1.96V/m occurs near the edges/over the wire windings of the circular coil as expected.

**Table S.1 Categorisation of neuronal subtypes in the cell cultures.**

| Excitatory neurons | Inhibitory neurons | Astrocytes | Other |
|---|---|---|---|
| 12.2% | 71.7% | 6% | 10.1% |

The percentages of excitatory (GAD65-67 negative) neurons, inhibitory (GAD65-67 positive) neurons, and astrocytes (GFAP positive) were quantified from two coverslips over two culture runs.
